# Concordance and incongruence in preclinical anxiety models: systematic review and meta-analyses

**DOI:** 10.1101/020701

**Authors:** Farhan Mohammad, Joses Ho, Jia Hern Woo, Chun Lei Lim, Dennis Jun Jie Poon, Bhumika Lamba, Adam Claridge-Chang

## Abstract

Rodent defense behavior assays have been widely used as preclinical models of anxiety to study possibly therapeutic anxiety-reducing interventions. However, some proposed anxiety-modulating factors—genes, drugs and stressors—have had discordant effects across different studies. To reconcile the effect sizes of purported anxiety factors, we conducted systematic review and meta-analyses of the literature on ten anxiety-linked interventions, as examined in the elevated plus maze, open field and light-dark box assays. Diazepam, 5-HT1A receptor gene knockout and overexpression, SERT gene knockout and overexpression, pain, restraint, social isolation, corticotropin-releasing hormone and Crhr1 were selected for review. Eight interventions had statistically significant effects on rodent anxiety, while Htr1a overexpression and Crh knockout did not. Evidence for publication bias was found in the diazepam, Htt knockout, and social isolation literatures. The Htr1a and Crhr1 results indicate a disconnect between preclinical science and clinical research. Furthermore, the meta-analytic data confirmed that genetic SERT anxiety effects were paradoxical in the context of the clinical use of SERT inhibitors to reduce anxiety.

**Highlights:**

- Meta-analysis shows eight rodent anxiety factors have at least moderate effects.
- Publication bias affects four of the anxiety interventions.
- Preclinical rodent anxiety results appear disconnected from clinical efforts.
- Serotonin transporter gene lesion effects are paradoxical with reuptake inhibitors clinical use.

## 1 Introduction

The anxiety disorders are among the costliest classes of mental disorders, with regard to both morbidity and economic cost (Baldwin et al., 2014; DiLuca and Olesen, 2014). Development of anxiety-reducing (anxiolytic) drugs has been a major focus of the pharmaceutical industry and academic neuropsychiatric research, though no new drug types have been adopted since the introduction of selective serotonin uptake inhibitors (SSRIs) and other antidepressants for the treatment of anxiety disorders (Griebel and Holmes, 2013; Tone, 2009). Anxiety research relies on similarities between human emotional behavior and behaviors in animals (Darwin, 1998), specifically rat and mouse (Prut and Belzung, 2003). While there are many rodent behavioral paradigms that aim to model anxious behavior, three **a**nxiety-**r**elated **de**fense **b**ehavior (ARDEB) assays that specifically aim to measure rodent anxiety have been widely adopted, also referred to as ‘approach-avoidance conflict tests’: the elevated plus maze (EPM), the light-dark box (LD) and the open field (OF), the first, second and fifth most widely used rodent anxiety assays, respectively (Griebel and Holmes, 2013). All three assays use an arena that contains a sheltered domain (e.g., the closed arms in EPM) and an exposed region. It is believed that an animal’s avoidance of the exposed portions of the chamber reports on anxiety-like brain states. The ARDEB assays are accepted as preclinical assays of anxiety disorders, by reference to classic studies that tested their predictive validity with panels of drugs known to have anxiety-modulating effects in humans (Crawley and Goodwin, 1980; Pellow et al., 1985; Simon et al., 1994).

Rodent research has been implicated in the largely frustrated efforts to develop new types of anxiolytics (Griebel and Holmes, 2013). The literature regarding defense behaviors is contradictory about the size and even the direction of many interventions that are proposed to be anxiolytic or anxiogenic (together ‘anxiotropic’) (Griebel and Holmes, 2013; Prut and Belzung, 2003). This is true even for some anxiety-related factors with major clinical relevance, such as the serotonin transporter (SERT/*Htt*), the target of the SSRIs. As with the assessment of clinical anxiety interventions (Baldwin et al., 2014), a solid preclinical evidence base is necessary to guide decisions about further research and therapeutic development (Vesterinen et al., 2014). To better understand the widespread discordance in rodent anxiety studies, we conducted a quantitative review of the effect of purported anxiety factors on rodent ARDEB. The primary aim of this study was to examine the relevance of these factors and to estimate the magnitude of their effects on rodent anxiety. A secondary goal of this analysis was to examine patterns in ARDEB factor evidence: gaps in the literature, the extent of standardization/heterogeneity and publication bias. Synthesizing the data on anxiety-targeted interventions might also assist in understanding why these assays have not led to new therapies. Once confirmed by meta-analysis, effective anxiotropic interventions can be adopted as benchmarks against which to validate new rodent assays and/or more tractable model animal species (e.g. *Drosophila* and zebrafish).

## 2 Materials and Methods

### 2.1 Literature review

We identified genes, drugs and environmental interventions that had been proposed to be involved in anxiety with a literature search of anxiety review articles. From two histories of anxiety research (Griebel and Holmes, 2013; Tone, 2009), a list of ten anxiotropic interventions were chosen to be included in the systematic review, either due to their clinical relevance (e.g., diazepam, Htt), their role as an example of a class of proposed anxiety-related factors (e.g., isolation), or their connection to possible forthcoming therapeutics (e.g. Crh). A systematic review was conducted to identify published articles addressing experimental outcomes in rodents from the EPM, OF, or LD assays for these interventions (Figure 1). The literature for each genetic, pharmacological or environmental intervention was identified by a search of PubMed and EMBASE using specific search phrases (Table 1). The selective serotonin reuptake inhibitors (SSRIs), which have clinical importance (Baldwin et al., 2014), a very large number of studies conducted on them (Griebel and Holmes, 2013), and controversial efficacy (Kirsch et al., 2008) are the subject of a separate meta-analytic study, currently in preparation.

### 2.2 Eligibility criteria and study selection

The search phrases in Table 1 were used to identify lists of studies. We exported the articles’ bibliographic data (including study ID, date of publication, title and abstract) of to a spreadsheet. Each article on this list was then reviewed at one or more of four levels of detail (title, abstract, full text and a detailed review of experimental design) to determine their eligibility for the review. Studies were required to be written in English and to have reported ARDEB in adult rats or mice. We required that each included study contain (1) primary behavior data from either an OF, EPM, or LD experiment for at least one of the interventions of interest, (2) suitable control data and (3) the relevant statistics (mean, standard error or standard deviation, and sample sizes of both control and intervention groups). Experiments that used combination treatments were excluded. Only studies in which adult rodents were assayed were included. For gene knockout and overexpression interventions, we included only experiments that used a lifetime loss of function throughout the entire animal. All eligible experiments from all eligible studies were included in the ten meta-analyses (Table 1).

### 2.3 Data items and extraction

The following data were collected from each of the included studies: authors, year of publication, figure and panel numbers, species, genotype, and mean, standard error of the mean and sample size (N) of each intervention and its related control group. Graphically presented data were extracted from Portable Document Format (PDF) files with the Measuring Tool in Adobe Acrobat Pro. All extracted data were checked by a second researcher. For values extracted from tables, the check consisted of ensuring the values were identical. For values extracted from graphical data (e.g. bar plots), the check consisted of a visual inspection to ensure that the extracted value matched the graphical data. Extraction discrepancies were reconciled by conference between the primary extractor and the researcher who identified the discrepancy.

**Table I.** **Summary of systematic reviews of anxiety-related interventions in mouse and rat.** The PubMed and Embase query phrases used to identify articles that might contain data relevant to the interventions and assays of interest are detailed. Title, abstract and full-text searches were performed to identify articles meeting the selected criteria.

| Intervention | Query phrase used in PubMed and Embase for study selection | Articles identified by phrase | Articles meeting meta-analysis criteria |
| --- | --- | --- | --- |
| **Diazepam** | (diazepam OR valium) AND anxiety AND (open field OR exploratory) AND (rodent OR rat OR rats OR mouse OR mice OR *Mus*) | 540 | 172 |
| ***Ht1a* knockout** | (serotonin1A receptor OR 5-HT1A receptor) AND knockout AND anxiety | 85 | 12 |
| ***Ht1a* over-expression** | (serotonin1A receptor OR 5-HT1A receptor) AND (overexpression OR over-expression OR overexpressing) AND anxiety | 13 | 3 |
| ***Htt* knockout** | (serotonin transporter) AND (knockout OR knockdown OR deletion OR antisense) AND anxiety AND (elevated plus maze OR open field OR light-dark) AND (rats OR rat OR mice OR mouse OR *Mus*) | 37 | 13 |
| ***Htt* over-expression** | serotonin transporter AND anxiety AND (elevated plus maze OR open field OR light-dark) AND (increased OR over-expression OR overexpressing OR transgenic) AND (rats OR rat OR mice OR mouse OR *Mus*) | 65 | 2 |
| ***Crh* knockout** | ((corticotropin-releasing) AND (stress-induced behaviors OR stress-related behaviors or Behavioral responses to stress OR Behavioral responses to stress)) AND (CRH-deficient mice OR lacking the CRH gene OR CRFko) | 12 | 2 |
| ***Crhr1* knockout** | (corticotropin releasing factor receptor 1-deficient mice) OR CRH1 receptor antisense oligodeoxynucleotide OR *Crhr1* null mutants OR Corticotropin-releasing hormone receptor antisense | 62 | 6 |
| **Pain** | (inflammatory pain OR neuropathic pain) AND anxiety AND (elevated plus maze OR open field OR light-dark) AND (rats OR rat OR mice OR mouse OR *Mus*) | 73 | 7 |
| **Restraint** | acute restraint AND anxiety AND (elevated plus maze OR open field OR light-dark) AND (rats OR rat OR mice OR mouse OR *Mus*) | 87 | 15 |
| **Isolation** | (social isolation OR single housing) AND anxiety AND (elevated plus maze OR plus-maze OR open field OR OFT OR light-dark) AND (rats OR rat OR mice OR mouse OR *Mus*) | 167 | 50 |

**Figure 1.**
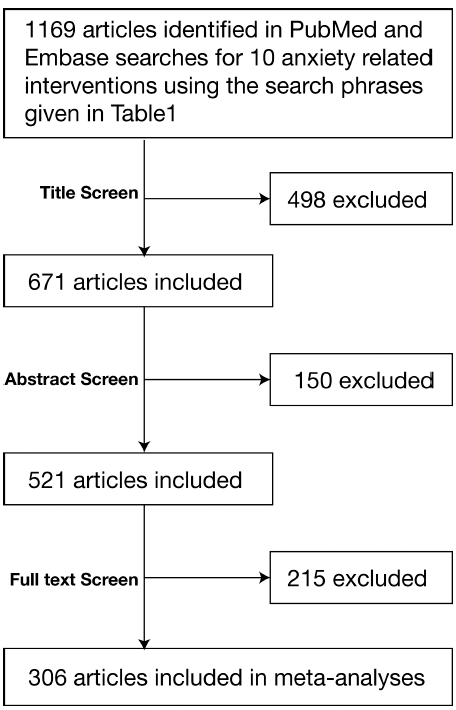
**Flow chart of the systematic literature review of 10 anxiotropic interventions.** The literature was reviewed in a four-stage process, starting with searches of the Pubmed and EMBASE databases that yielded 1169 articles, followed by three screens of increasing detail, reviewing the article title, abstract, and full text for experimental design. A total of 306 articles were used in the meta-analysis. Further details are given in Table 1 and the Methods section.

### 2.4 Summary measures

The following behavioral metrics were extracted from the articles: in OF studies, percent or total time spent at the center; in EPM studies, percent or total time spent on the open arm; in LD studies, percent or total time spent in the bright area. To synthesize these time-based metrics from the three assays, all estimates were standardized to Hedges’ *g*, a preferred variant of Cohen’s d that uses the pooled standard deviation and is corrected for bias using Hedges’ method (Borenstein et al., 2011; Cumming, 2012). The conventional adjectives to describe effect size—trivial, small, moderate, large—are used for effect sizes *g* < 0.2, *g* < 0.5, *g* < 0.8 and *g* > 0.8 SD respectively (Cumming, 2012).

### 2.5 Synthesis of results

Meta-analyses of experimental outcomes, including the calculation of weighted mean effect sizes (Hedges’ *g*), 95% confidence intervals, I^2^ heterogeneity values, and P values using the random effects model, `were performed with the metafor package in R` (http://CRAN.R-project.org/package=metafor) (Viechtbauer, 2010). All error bars in forest plots are 95% confidence intervals; forest plots were generated with custom R scripts.

### 2.6 Assessment of bias across studies

Publication bias was assessed with funnel plots and Egger’s linear regression test of funnel plot asymmetry (Egger et al., 1997). The standard normal deviate (Hedges’ *g* / standard error) for each study was regressed against the study’s precision (1 / standard error) using the “lm” function in R (http://www.R-project.org/). For studies that showed publication bias (P-value ≤ 0.05), the trim-and-fill method (Duval and Tweedie, 2000) was employed to estimate the effects of publication bias on the effect size estimate. Funnel plots and trim-and-fill adjustments were performed with the ‘metafor’ package in R (Viechtbauer, 2010).

## 3 Results

### 3.1 Review selection criteria identified 306 eligible articles

The flow-chart in Figure 1 summarizes the study selection process. In total, 1169 articles were identified by the initial search in PubMed and EMBASE databases. According to the selection criteria described above, 498 studies were excluded based on their titles and a further 150 were excluded based on their abstracts. The full text of the remaining 521 articles were screened for criteria related to experimental paradigm, methods, and relevant variables, resulting in the exclusion of a further 215 studies. A total of 306 articles were considered eligible for inclusion in the review.

### 3.2 Characteristics of included experiments

The characteristics of all included studies are given in Supplementary Table 1. In brief, 582 experiments from 306 studies comprising 411 EPM experiments, 84 OF experiments and 87 LD experiments were identified. Studies were published between 1985 and 2015 and included data from 318 experiments conducted on mice and 264 experiments on rats. Studies reported 515 experiments conducted on male animals, 29 on female, 35 on mixed and 3 experiments with no gender information reported. ARDEB studies of diazepam used a median dosage of 1 mg/kg, with minimum and maximum dosages of 0.01 mg/kg and 20 mg/kg respectively, a dose range is similar to or higher than commonly used by patients.

### 3.3 Heterogeneity

Statistically significant heterogeneity was found in (8/10) of the metaanalyses. Only two meta-analyses had high heterogeneity, I^2^ > 75%: *Htr1a* overexpression, and physical restraint (Higgins et al., 2003).

**Figure 2.**
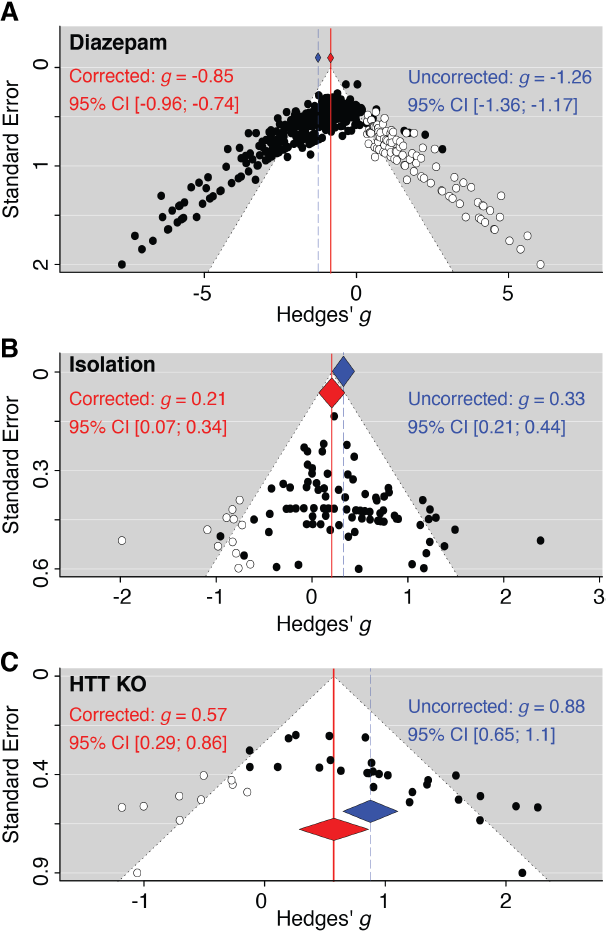
**Funnel plots of three meta-analyses with evidence for publication bias.** Where at least ten experiments were available for meta-analysis, the effect sizes (Hedges’ *g*) of the experiments are plotted against their respective standard errors. Points on each plot represent individual experiments. The triangle bounded by dotted lines indicates the area where 95% of studies are expected to fall, in the absence of both publication bias and study heterogeneity. Shown here are funnel plots for experiments on (A) diazepam, (B) social isolation, --and (C) *Htt* knockout.

Three of the meta-analyses, pain and *Htt* knockouts and diazepam, had moderate heterogeneity (50% < I^2^ < 75%). Five meta-analyses had low heterogeneity (I^2^ < 50%). As most of these syntheses contained data from more than one assay type, it is encouraging that half had low or moderate heterogeneity, and outcome that is compatible with the idea that the three ARDEB assays are testing similar aspects of rodent anxiety.

### 3.4 Substantial publication bias in four anxiety factors

Censorship of non-statistically significant experimental results and selective publication of statistically significant ‘positive’ results can cause a literature (and meta-analysis thereof) to overstate effect sizes. This effect, termed ‘publication bias,’ has a profound influence on the literature on rodent models of stroke, and may affect other animal models (Sena et al., 2010). Publication bias in the ARDEB literature was assessed for the six meta-analyses that had at least 20 experiments (Table 2) (Sterne et al., 2011). Funnel plots of these data showed pronounced asymmetry (Figure 2), which pointed to publication bias in these literatures (Sterne et al., 2011). Egger’s asymmetry test indicated that four of these literatures showed statistically significant bias (Table 2). For the biased data sets, we applied trim-and-fill adjustment to estimate the number of hypothesized missing studies and to correct the bias (Duval and Tweedie, 2000) (Figure 2). These data support the idea that the literatures of diazepam, *Htt* knockout, social isolation and restraint effects on ARDEB are strongly affected by publication bias.

**Table 2.**
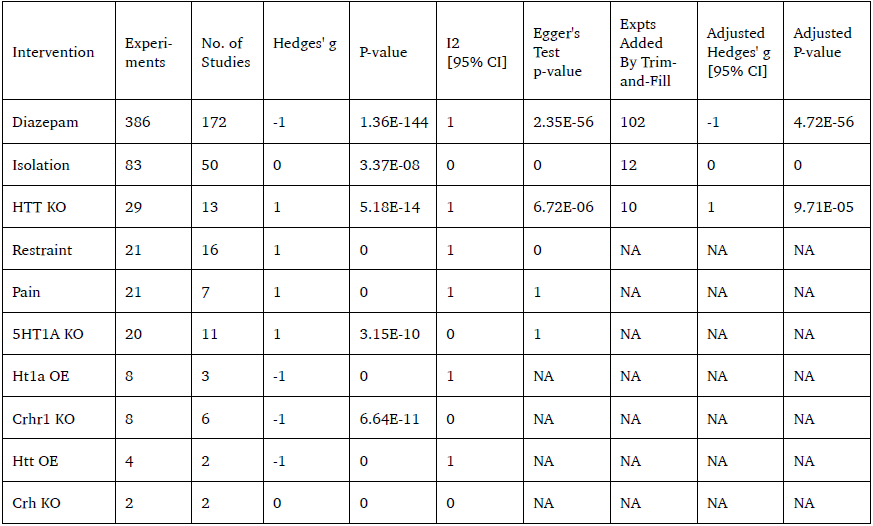
**Results of Egger’s linear regression test for funnel plot asymmetry across six meta-analyses** Where at least twenty experiments were available for meta-analysis, Egger’s linear regression test for funnel plot asymmetry was performed. For each meta-analysis, the number of included studies, the vertical intercept of the linear regression, and the P-values of Egger’s test are listed.

**Figure 3.**
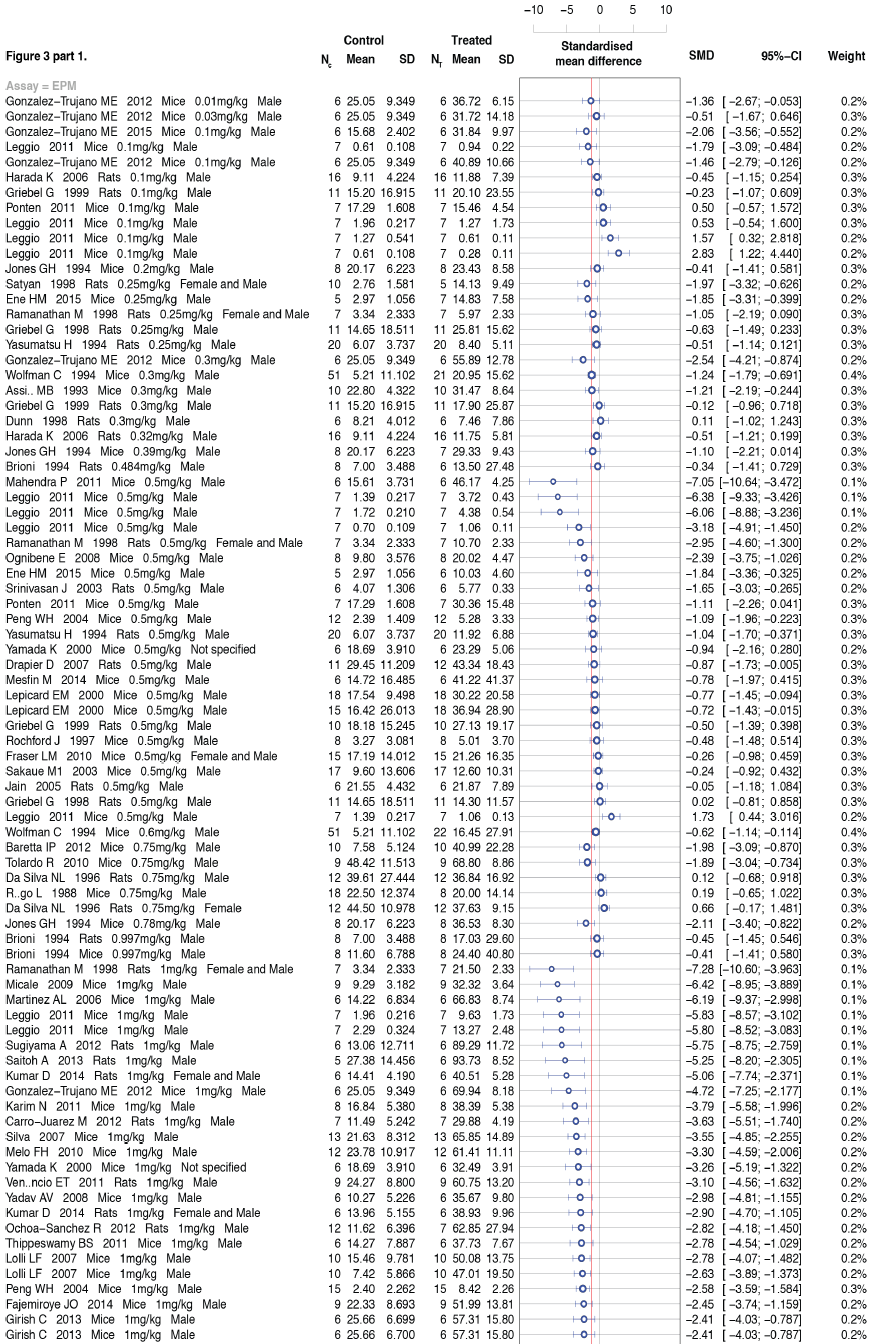
**Meta-analysis of diazepam on rodent anxiety related behavior.** Meta-analysis of rodent diazepam effect sizes, shown as a forest plot of standardized effect sizes (Hedges’ *g*). The meta-analysis is sub-grouped by animal species. Error bars indicate the 95% confidence intervals of standardized mean difference. The weighted average mean effect size of subgroups and all studies is represented by the central vertices of a red diamond; the outer vertices indicate the 95% confidence intervals. Control and treatment samples sizes are given in the columns listed as N_C_ and N_T_ respectively.

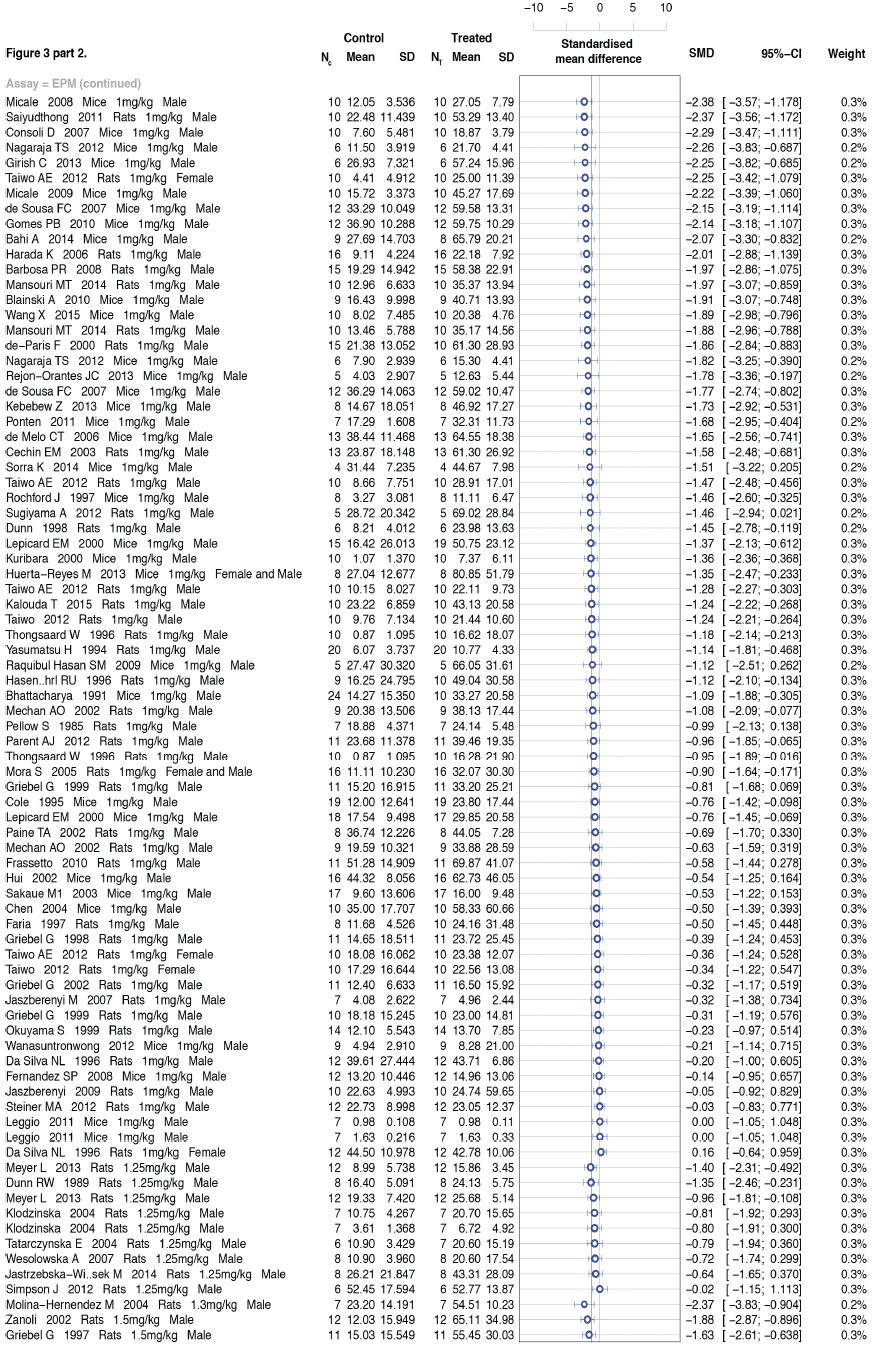

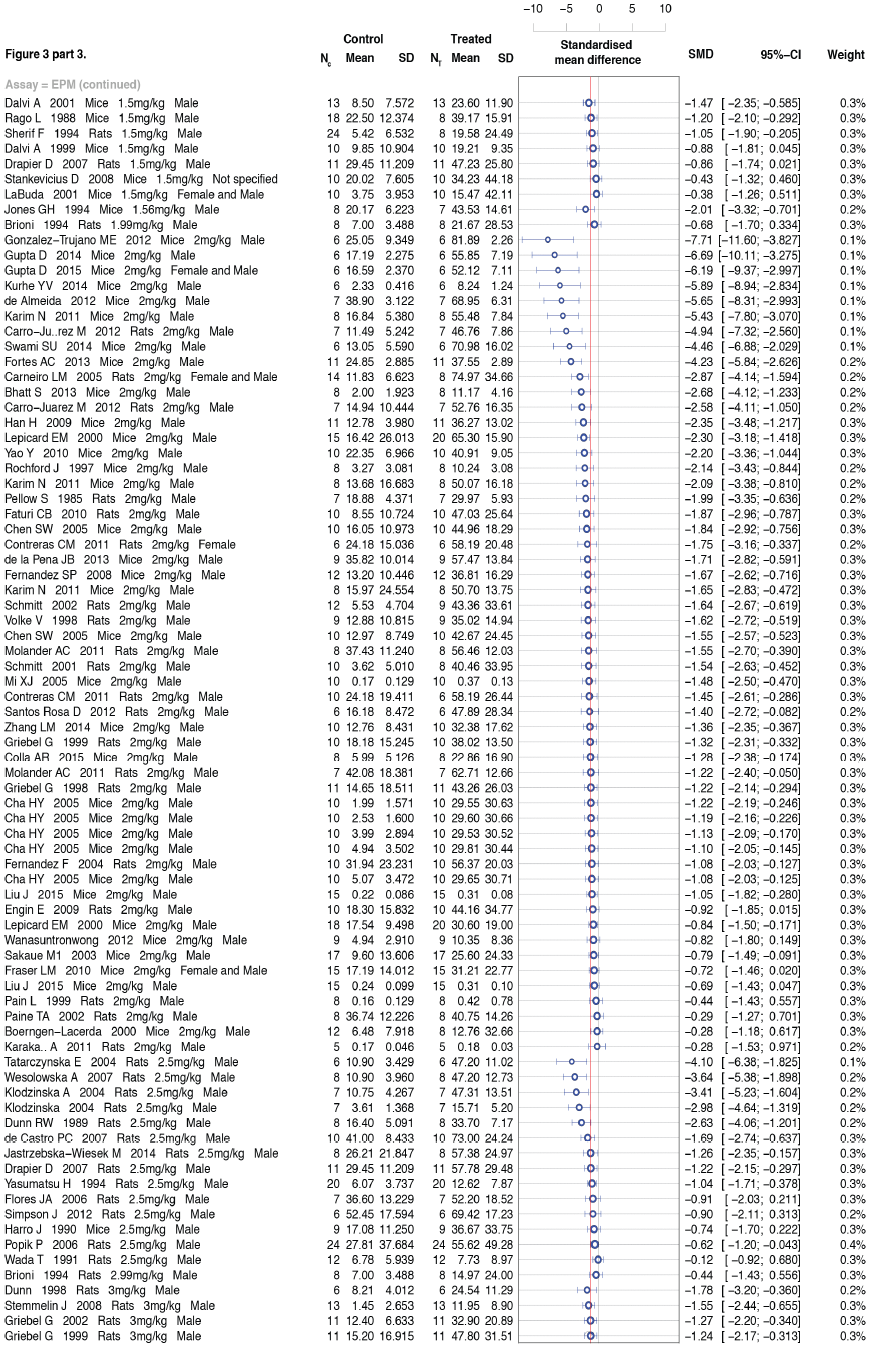

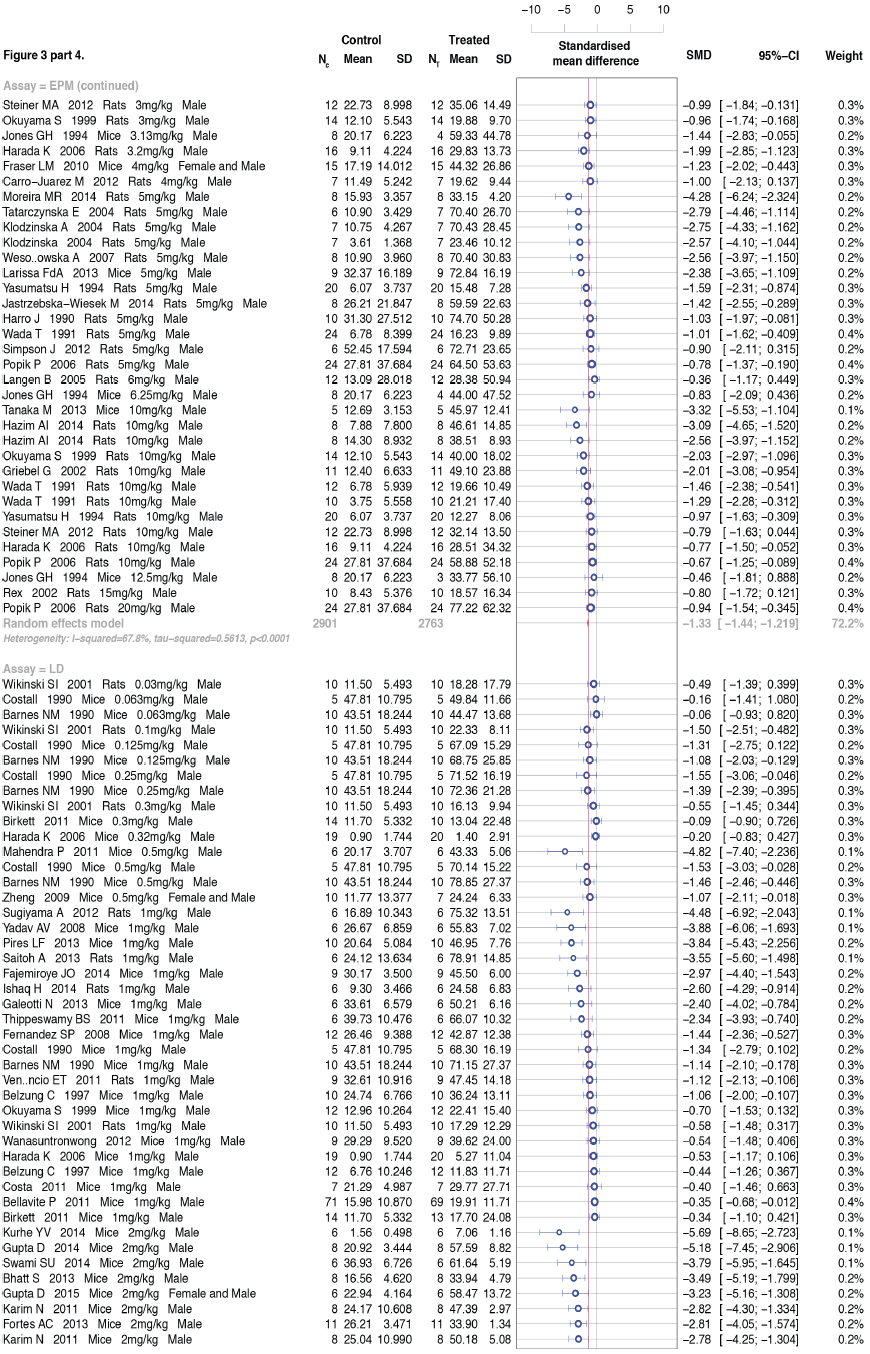

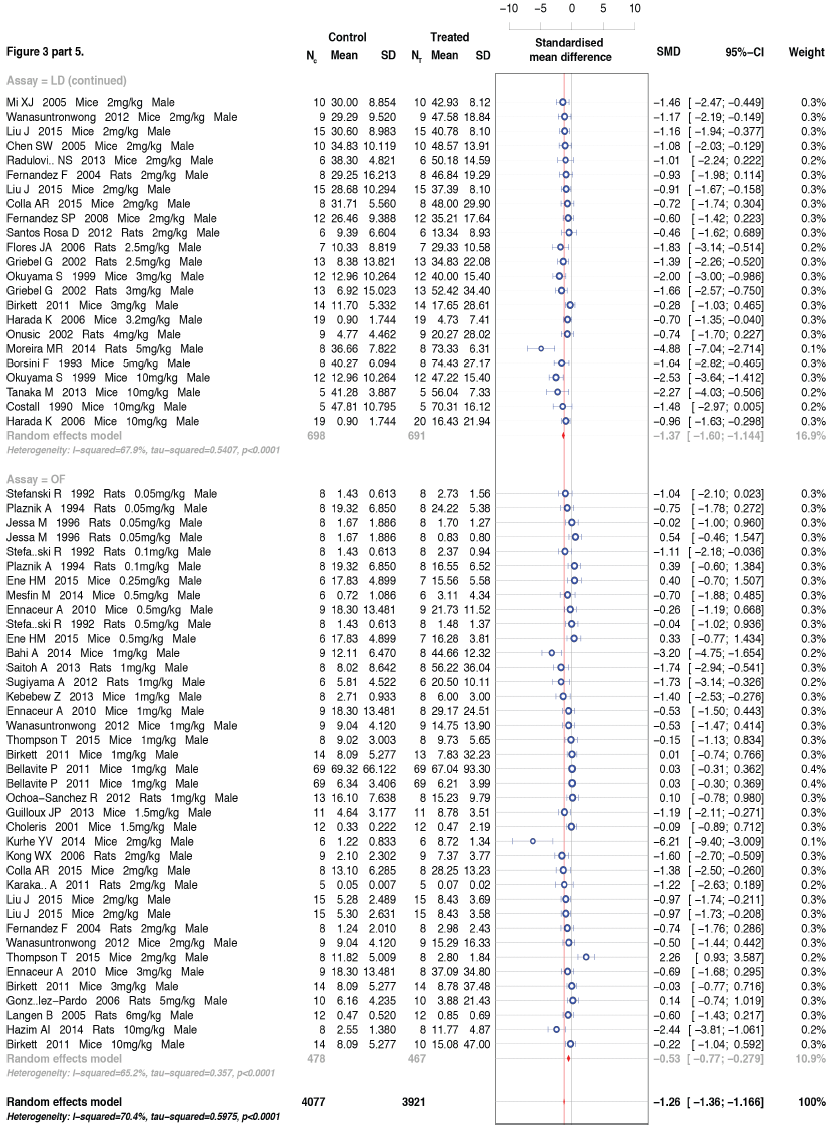

### 3.5 Diazepam reduces anxiety-related defense behaviors

Diazepam is an important minor tranquilizer that was used for decades as the first line of treatment for anxiety disorders (Tone, 2009) and, along with other benzodiazepines, is still used extensively to control anxiety (Baldwin et al., 2014). Recent clinical meta-analysis studies have found support for the efficacy of benzodiazepines in the short-term treatment of anxiety disorders (Baldwin et al., 2014). However, a review of diazepam effects in open field studies revealed widespread disagreement between with 29 studies supporting an anxiolytic effect and 23 supporting either an anxiogenic effect or no effect (Prut and Belzung, 2003). We reviewed the available literature on diazepam for the three major rodent ARDEB assays: EPM, OF and LD. This review identified 172 articles containing relevant data (Assie et al., 1993; Bahi et al., 2014; Barbosa et al., 2008; Baretta et al., 2012; Barnes et al., 1990; Bellavite et al., 2011; Belzung and Agmo, 1997; Bhatt et al., 2013; Bhattacharya and Mitra, 1991; Birkett et al., 2011; Blainski et al., 2010; Borsini et al., 1993; Brioni et al., 1994; Carneiro et al., 2005; Carro-Juarez et al., 2012; Cechin et al., 2003; Cha et al., 2005; Chen et al., 2004; 2005; Choleris et al., 2001; Cole and Rodgers, 1995; Colla et al., 2015; Consoli et al., 2007; Contreras et al., 2011; Costa et al., 2011; Costall et al., 1990; Da Silva et al., 1996; Dalvi and Rodgers, 2001; 1999; de A Vieira et al., 2013; de Almeida et al., 2012; de Castro et al., 2007; de Melo et al., 2006; de Sousa et al., 2007; de-Paris et al., 2000; Drapier et al., 2007; R. W. Dunn et al., 1989; R. W. Dunn et al., 1998; Ene et al., 2015; Engin et al., 2009; Ennaceur et al., 2010; Fajemiroye et al., 2014; Faria et al., 1997; Faturi et al., 2010; F. Fernandez et al., 2004; S. P Fernandez et al., 2008; Flores et al., 2006; Fortes et al., 2013; Fraser et al., 2010; Frassetto et al., 2010; Galeotti et al., 2013; Girish et al., 2013; Gomes et al., 2010; Ma Eva Gonzalez-Trujano et al., 2006; Maria Eva Gonzalez-Trujano et al., 2015; González-Pardo et al., 2006; Griebel et al., 1998; 1997; 1999a; 1999b; 2002; Guilloux et al., 2013; Gupta et al., 2014; 2015; Han et al., 2009; Harada et al., 2006; Hasenohrl et al., 1996; Hazim et al., 2014; Huerta-Reyes et al., 2013; Hui et al., 2002; Ishaq, 2014; N. S. Jain et al., 2005; Jastrzebska-Wiesek et al., 2014; Jászberényi et al., 2009; Jászberényi et al., 2007; Jessa et al., 1996; Jones et al., 1994; Kalouda and Pitsikas, 2015; Karakas et al., 2011; Karim et al., 2011; Kebebew and Shibeshi, 2013; Klodzinska et al., 2004a; 2004b; Kong et al., 2006; Kumar and Bhat, 2014; Kurhe et al., 2014; Kuribara et al., 2000; la Pena et al., 2013; LaBuda and Fuchs, 2001; Langen et al., 2005; Leggio et al., 2011; Lepicard et al., 2000; Jie Liu et al., 2015; Lolli et al., 2007; Mahendra and Bisht, 2011; Mansouri et al., 2014; Martinez et al., 2006; Mechan et al., 2002; Melo et al., 2010; Mesfin et al., 2014; Meyer et al., 2013; Mi et al., 2005; Micale et al., 2009; 2008; Molander et al., 2011; Molina-Hernandez et al., 2004; Mora et al., 2005; Moreira et al., 2014; Nagaraja et al., 2012; Ochoa-Sanchez et al., 2012; Ognibene et al., 2008; Okuyama et al., 1999; Onusic et al., 2002; Pain et al., 1999; Paine et al., 2002; Parent et al., 2012; Pellow et al., 1985; Peng et al., 2004; Pires et al., 2013; Plaznik et al., 1994; Ponten et al., 2011; Popik et al., 2006; Radulovic et al., 2013; Rago et al., 1988; Ramanathan et al., 1998; Raquibul Hasan et al., 2009; Rejon-Orantes et al., 2013; Rex et al., 2002; Rochford et al., 1997; Saiyudthong and Marsden, 2011; Sakaue et al., 2003; Santos Rosa et al., 2012; Satyan et al., 1998; Schmitt et al., 2002; 2001; Sherif et al., 1994; Silva et al., 2007; Simpson and Kelly, 2012; Sorra et al., 2014; Srinivasan et al., 2003; Stankevicius et al., 2008; Stefanski et al., 1992; Steiner et al., 2012; Stemmelin et al., 2008; Sugiyama et al., 2012; Swami et al., 2014; Taiwo et al., 2012; Tanaka et al., 2013; Tatarczynska et al., 2004; Thippeswamy et al., 2011; Thompson et al., 2015; Thongsaard et al., 1996; Tolardo et al., 2010; Varty et al., 2002; Venancio et al., 2011; Volke et al., 1998; Wada and Fukuda, 1991; Wanasuntronwong et al., 2012; Wang et al., 2015; Wesolowska and Nikiforuk, 2007; Wikinski et al., 2001; Wolfman et al., 1994; Yadav et al., 2008; Yamada et al., 2000; Yao et al., 2010; Yasumatsu et al., 1994; Zanoli et al., 2002; L.-M. Zhang et al., 2014; Zheng et al., 2009). Calculation of an average Hedges’ g (Cumming, 2012) for the 386 experiments contained therein indicated that diazepam had a very large effect on ARDEB, with a -1.26 *g* [95CI -1.36, -1.17] reduction compared with untreated control animals (Figure 3, Table 2). However, as Egger’s regression indicated the source literature was affected by publication bias, trim-and-fill correction indicated a smaller—though still large—effect of -0.85 *g* [95CI -0.74, -0.96]. The meta-analysis had a moderate level of heterogeneity (I^2^ = 70.4%). Subgroup analysis of assay types suggest that assays were not major source of heterogeneity (Figure 3). Subgroup analysis of the assay types showed that EPM and LD yielded very similar diazepam effect sizes (g = -1.33 and -1.37 respectively, Figure 3). However, the diazepam effect size in OF was found to be substantially smaller (g = -0.53, Figure 3). Additional subgroup analyses of treatment duration variables, dosages and route of treatments revealed that these factors were also only minor sources of heterogeneity (data not shown), indicating that laboratory, dosage, strain and other possible sources of experimental variation likely play a role. As diazepam is universally accepted to be an effective anxiolytic, and it has a robust (bias-corrected) effect in the rodent ARDEB assays, this meta-analytic result verifies the validity of the defense behavior tests.

**Figure 4.**
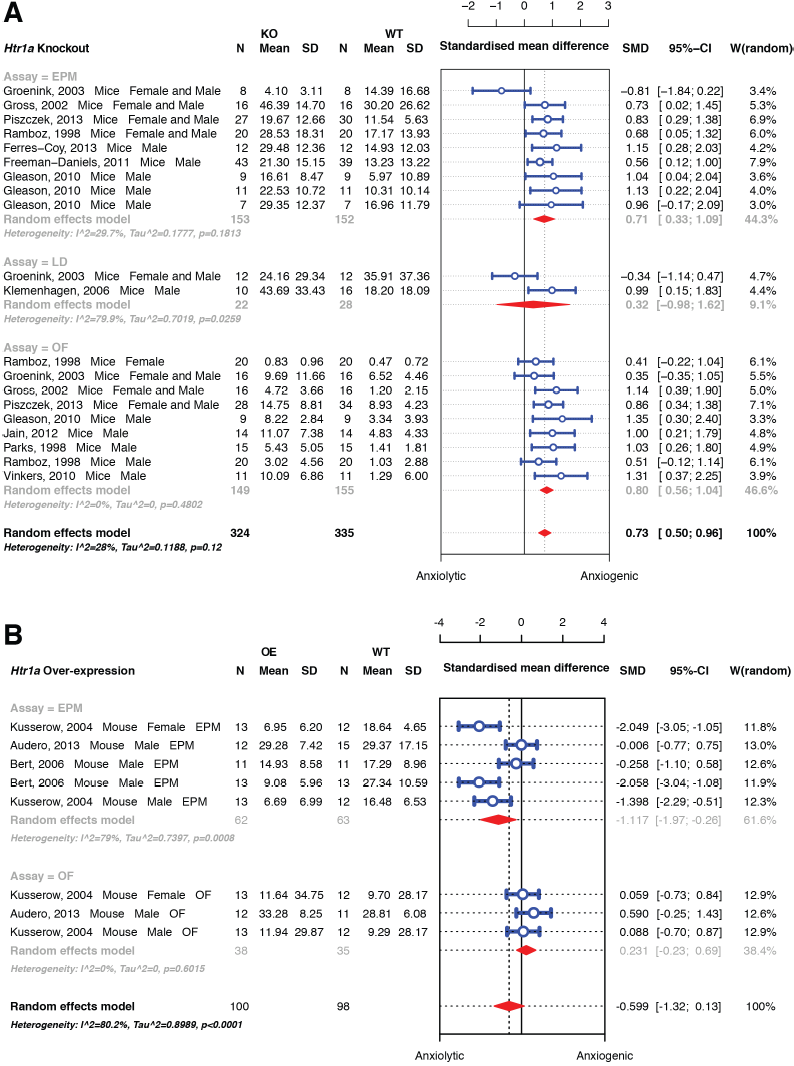
**Meta-analyses of serotonin receptor 1A interventions on rodent anxiety-related behaviors.** Meta-analysis of effect sizes of serotonin-targeted interventions is shown as a forest plot of standardized effect sizes (Hedges’ *g*). Error bars indicate the 95% confidence intervals of *g*. The weighted average mean effect size of all studies is represented by the central vertices of a red diamond; the outer vertices indicate the 95% confidence intervals. Control and treatment samples sizes (N_C_, N_T_) and the assay types of the studies are given; elevated plus maze (EPM), open field (OF) and light-dark box (LD). Effects of: A. Serotonin receptor gene Htrla knockout models. B. Htrla overexpression.

**Figure 5.**
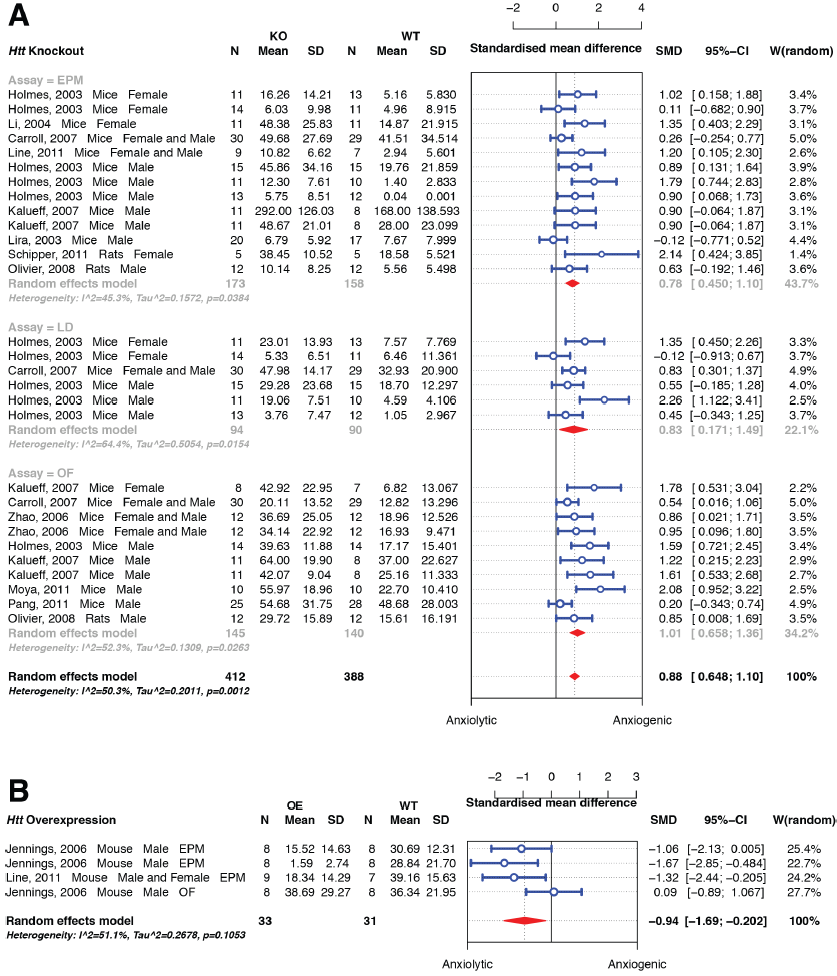
**Meta-analyses of serotonin transporter interventions on rodent anxiety-related behaviors.** Meta-analysis of effect sizes of serotonin-targeted interventions is shown as a forest plot of standardized effect sizes (Hedges’ *g*). Error bars indicate the 95% confidence intervals of *g*. The weighted average mean effect size of all studies is represented by the central vertices of a red diamond; the outer vertices indicate the 95% confidence intervals. Control and treatment samples sizes (N_C_, N_T_) and the assay types of the studies are given; elevated plus maze (EPM), open field (OF) and light-dark box (LD). Effects of: A. Serotonin transporter gene (Htt) knockout models B. Htt overexpression models.

**Figure 6.**
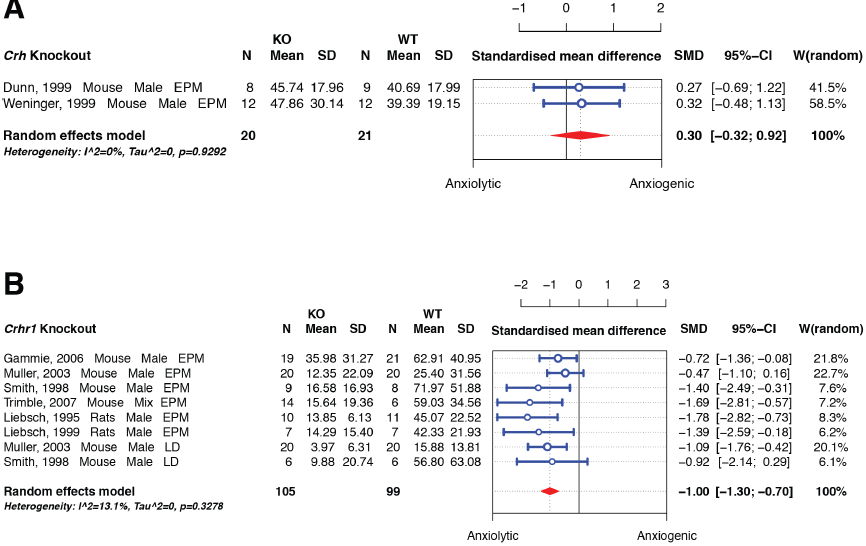
**Meta-analyses of the effects of stress signaling genes on anxiety-related behaviors.** Meta-analysis of effect sizes of stress signaling genes, shown as a forest plot of standardized effect sizes (Hedges’ *g*). Error bars indicate the 95% confidence intervals of *g*. The weighted average mean effect size of all studies is represented by the central vertices of a red diamond; the outer vertices indicate the 95% confidence intervals. Control and treatment samples sizes (N_C_, N_T_) and the assay types of the studies are given; elevated plus maze (EPM), open field (OF) and light-dark box (LD). Effects of: A. Crh gene knockout models. B. Crhr1 gene knockout models.

### 3.6 5-HT1A receptor function influences ARDEB

Following negative publicity regarding the adverse effects of benzodiazepines (Tone, 2009), pharmaceutical companies focused on the serotonergic system (Griebel and Holmes, 2013). Of the fourteen mammalian serotonergic receptors, much investigation has centered on the serotonin receptor 5-HT1A and its proposed influence on anxiety disorders and depression (Samuels et al., 2014). More than 1200 articles describe experiments connecting 5-HT1A agonism with rodent anxiety (Griebel and Holmes, 2013). However, a substantial proportion of those articles reported that 5-HT1A agonists or knockout of the *Htr1a* gene either produced no effect on anxiety or an effect that was opposite to the receptor’s proposed mode of action (Griebel and Holmes,2013). We systematically reviewed the literature on gene manipulations of *Htrla* and identified 11 knockout articles (Ferrés-Coy et al., 2013; Freeman-Daniels et al., 2011; Gleason et al., 2010; Groenink et al., 2003; Gross et al., 2002; A. Jain et al., 2012; Klemenhagen et al., 2006; Parks et al., 1998; Piszczek et al., 2013; Ramboz et al., 1998; Vinkers et al., 2010). Meta-analysis of the knockout data revealed that removal of *Htr1a* produced a moderate increase (Hedges’ *g* = 0.73 [95CI 0.50, 0.96], P = 3.5 × 10^−10^) in ARDEB phenotypes (Figure 4A). The three studies of *Htr1a* overexpression found by the review (Audero et al., 2013; Bert et al., 2006; Kusserow et al., 2004) indicated that this intervention moderately decreased ARDEB (*g* = −0.6 [95CI −1.3, 0.13], P = 0.11; Figure 4B). The cumulative sample size of *Htr1a* overexpression is quite substantial (N = 100, 98), though the moderate effect size observed was not statistically significant and had high heterogeneity (I^2^ = 80%). These results confirm that *Htr1a* function has a moderate effect on rodent anxiety.

### 3.7 Anxiotropic effects of the serotonin transporter

The serotonin transporter (SERT) is the target for the selective serotonin reuptake inhibitors (SSRIs), a class of drugs used to treat depression and anxiety (Baldwin et al., 2014). Meta-analysis of thirteen knockout studies (Carroll et al., 2007; Holmes et al., 2003a; Holmes et al., 2003b; Kalueff et al., 2007a; Kalueff et al., 2007b; Li et al., 2004; Line et al., 2011; Lira et al., 2003; Moya et al., 2011; Olivier et al., 2008; Pang et al., 2011; Schipper et al., 2011; Zhao et al., 2006) revealed a large anxiogenic effect (*g* = 0.88 [95CI 1.26, 0.23], P = 5.2 × 10^−14^; Figure 5A) produced by knocking out the SERT gene, *Htt*. However, a funnel plot and Egger’s regression revealed a pronounced bias in reported effect sizes (Egger’s test P = 6.7 × 10^−6^, Table 2). Trim-and-fill adjustment added ten imputed data points to the left segment of the funnel plot, lowering the effect size to *g* = 0.57 [95CI 0.29, 0.86], a moderate effect. Only two articles studying the effect of *Htt* overexpression on ARDEB were found (Jennings et al., 2006; Line et al., 2011). Meta-analysis revealed a large anxiolytic effect (*g* = −0.94 [95CI −1.69, −0.20], P = 0.013; Figure 5B) in EPM and OF assays (no LD articles were found). The transporter gene knockout and overexpression effects clearly connect *Htt* function to rodent anxiety. However, the direction of effects is the opposite of what would be expected from the clinical application of SERT inhibitors, given that SSRI reduction of SERT function is believed to have a therapeutic, anxiety-reducing effect.

**Figure 7.**
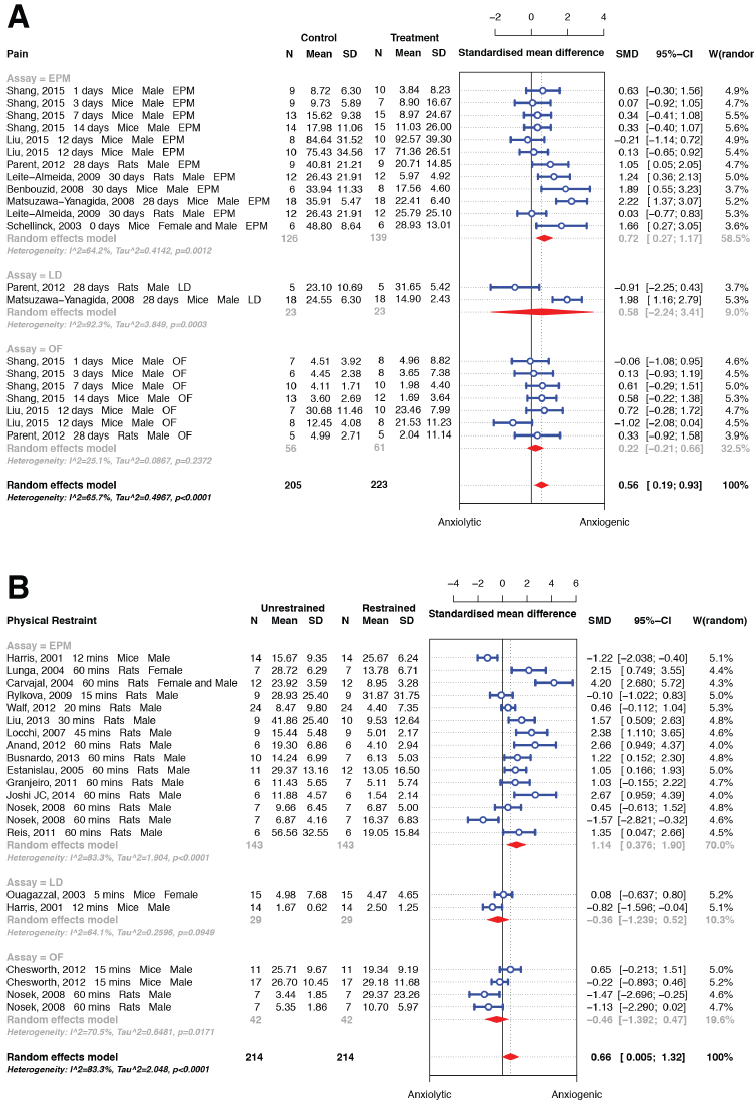
**Meta-analyses of experiments on the stress-anxiety relationship in rodents.** Meta-analysis of effect sizes of stress-anxiety interventions, shown as a forest plot of standardized effect sizes (Hedges’ *g*). Error bars indicate the 95% confidence intervals of *g*. The weighted average mean effect size of all studies is represented by the central vertices of a red diamond; the outer vertices indicate the 95% confidence intervals. Control and treatment samples sizes (N_C_, N_T_) and the assay types of the studies are given; elevated plus maze (EPM), open field (OF) and light-dark box (LD). Effects of: A. Acute pain. B. Restraint stress (immobilization). C. Social isolation.

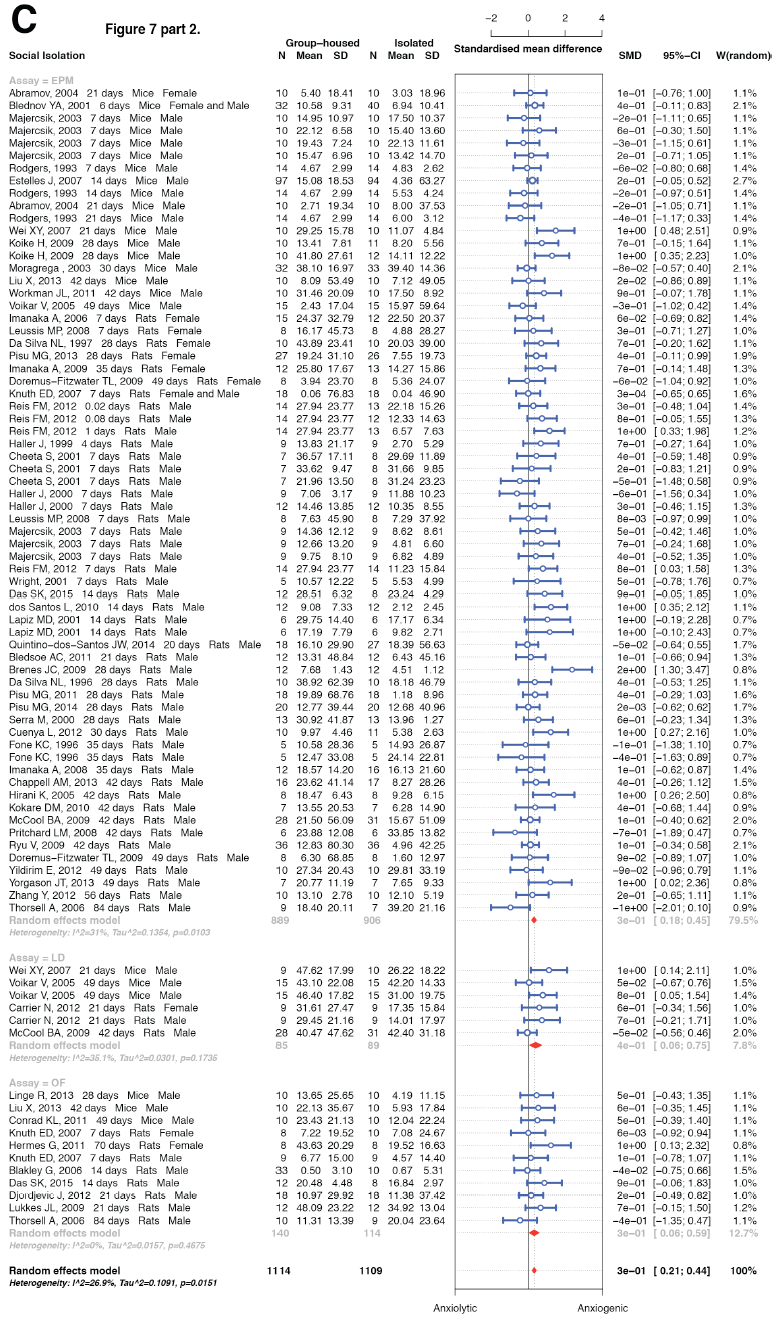

**Figure 8.**
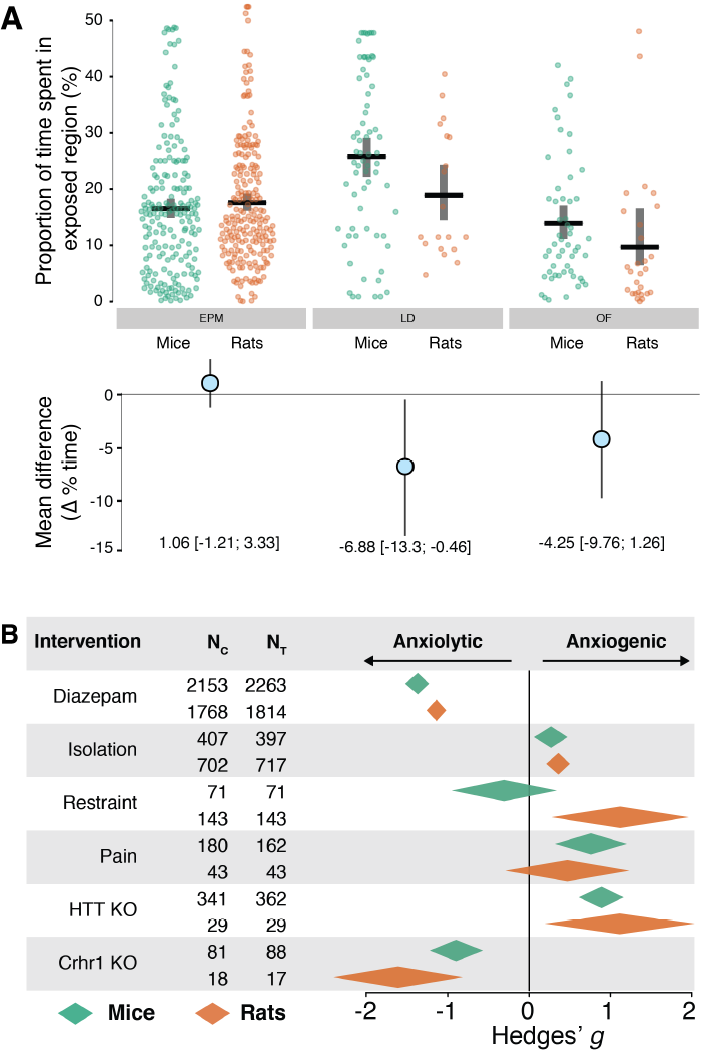
**Species differences in ARDEB.** A. Contrasts of mice and rats naive defence behaviors in three different assays. Upper panel shows the means of proportion of time spent in the exposed region, categorized by assay type and species. Each point is the mean value of an experiment. The lower panel shows the contrast means and confidence intervals (mean difference of the percent time spent in exposed region). B. The weighted mean effect sizes of six interventions subgrouped into mice and rats. Color indicates species (green = mice, orange = rats). Each mean effect size is represented by the central vertices of diamond; the outer vertices indicate the 95% confidence interval. The horizontal axis is Hedges’ *g*, the standard deviation change relative to control animals. N_C_ and N_T_ indicate control and treatment animal sample sizes respectively.

### 3.8 The effect of acute pain on rodent anxiety

Environmental stressors have physiological effects on animals that promote the anxiety-like state (van Praag, 2003). To survey a range of stress modalities we selected acute pain, bodily restraint and social isolation for review; all three have been found to promote anxiety in humans (Sherif and Oreland, 1995). The systematic review identified seven papers measuring the effect of acute pain on ARDEB (Benbouzid et al., 2008; Leite-Almeida et al., 2012; Yan Liu et al., 2015; Matsuzawa-Yanagida et al., 2008; Parent et al., 2012; Schellinck et al., 2003; Shang et al., 2014). Meta-analysis of the 21 experiments therein indicated a moderate anxiogenic effect (*g* = 0.56 [95CI 0.19, 0.93], P = 2.9 × 10^−3^; Figure 6A).

### 3.9 The effect of bodily restraint on rodent anxiety

Review of 16 studies of rodent bodily restraint (Anand et al., 2012; Busnardo et al., 2013; Carvajal et al., 2004; Chesworth et al., 2012; Estanislau and Morato, 2005; Granjeiro et al., 2011; Harris et al., 2001; Joshi et al., 2014; Jing Liu et al., 2011; Locchi et al., 2008; Lunga and Herbert, 2004; Nosek et al., 2008; Ouagazzal et al., 2003; D. G. Reis et al., 2011; Rylkova et al., 2009; Walf and Frye, 2012) containing 21 experiments indicated that it had an overall moderate anxiogenic effect in EPM and OF assays (*g* = 0.70 [ 95% CI 0.82 – 1.32], P = 0.027; Figure 6B). The restraint meta-analysis had a high level of heterogeneity, I^2^ = 89%; a subgroup analysis by assay type revealed that the different assays were not the source of this variability (data not shown).

### 3.10 Social isolation has a small effect on defense behaviors

Systematic review identified 50 articles on social isolation and ARDEB (Abramov et al., 2004; Blakley and Pohorecky, 2006; Blednov et al.,2001; Bledsoe et al., 2011; Brenes et al., 2009; Carrier and Kabbaj, 2012; Chappell et al., 2013; Cheeta et al., 2001; Conrad et al., 2011; Cuenya et al., 2012; Da Silva et al., 1996; Das et al., 2014; Djordjevic et al., 2012; Doremus-Fitzwater et al., 2009; Estelles et al., 2007; Fone et al., 1996; Haller and Halasz, 1999; Haller et al., 2000; Hermes et al., 2011; Hirani et al., 2005; Imanaka et al., 2008; Imanaka et al., 2006; Knuth and Etgen, 2007; Koike et al., 2009; Kokare et al., 2010; Lapiz et al., 2001; Leussis and Andersen, 2008; Linge et al., 2013; Xiao Liu et al., 2013; Lukkes et al., 2009; Majercsik et al., 2003; McCool and Chappell, 2009; Moragrega et al., 2003; Pisu et al., 2013; Pisu et al., 2011; Pritchard et al., 2008; Quintino-dos-Santos et al., 2014; F. M. C. V Reis et al., 2012; Rodgers and Cole, 1993; Ryu et al., 2009; Santos et al., 2010; Serra et al., 2000; Thorsell et al., 2006; Voikar et al., 2005; Wei et al., 2007; Workman et al., 2011; Wright and Ingenito, 2001; Yildirim et al., 2012; Yorgason et al., 2013; Y. Zhang et al., 2012). Meta-analysis revealed a small anxiogenic effect (*g* = 0.33 [95CI 0.21, 0.44], P = 3.4 × 10^−8^; Figure 6C), but this included likely publication bias; the trim-and-fill method corrected the anxiogenic effect to only 0.21 *g* [95CI 0.07, 0.34], P = 3.1 × 10^−3^) a very small anxiotropic effect (Figure 2B). It appears that, unlike the physical stressors, social isolation has only a modest influence on the ARDEB assays.

**Figure 9.**
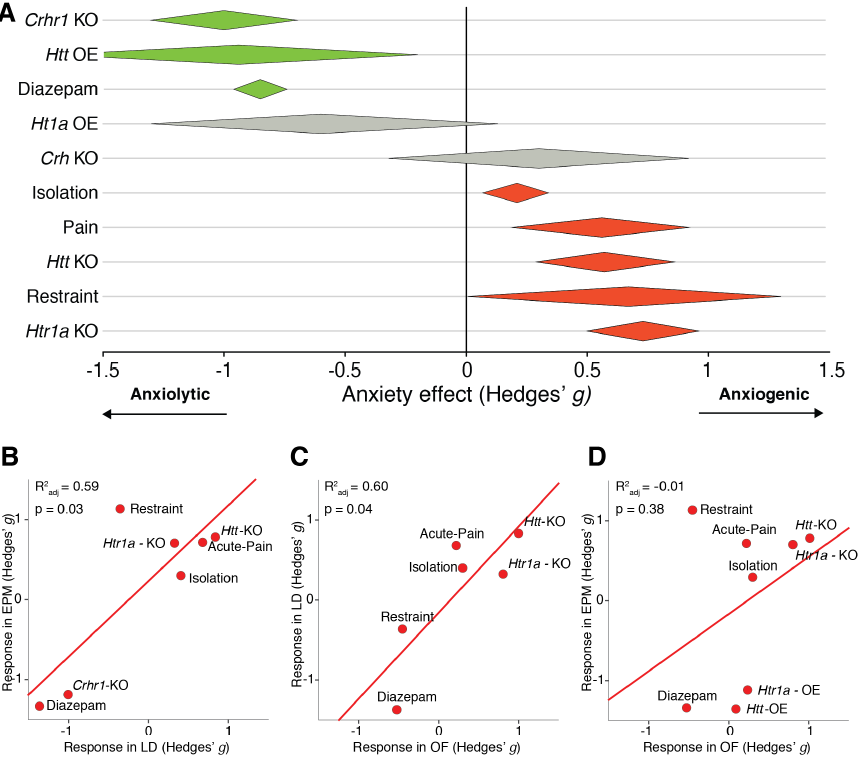
**Summary effect sizes of all meta-analyses.** A. The weighted mean effect sizes of all 10 interventions are shown here. Color indicates direction (green = anxiolytic, red = anxiogenic) and statistical significance (grey = statistically non-significant). The diamonds for the diazepam, social isolation, and Htt KO meta-analyses represent the summary effect sizes after trim-and-fill bias correction. Each mean effect size is represented by the central vertices of a diamond; the outer vertices indicate the 95% confidence intervals. The horizontal axis is Hedges’ *g*, the standard deviation change relative to control animals. B. Method comparison of LD and EPM shows that the two methods report ARDEB changes with 59% concordance. C. Method comparison of OF and LD shows that the two methods reports ARDEB changes with 60% concordance. D. Method comparion of OF and EPM shows that there is no concordance between the two methods.

### 3.11 Crh gene knockout has a modest effect on rodent ARDEB

Several neuropeptide-related genes involved in stress signaling have been linked to anxiety, notably the peptide, corticotropin-releasing hormone (CRH; also known as corticotropin-releasing factor) (Kormos and Gaszner, 2013) and its receptor, CRHR1. Two studies that examined the effects of *Crh* knockouts on ARDEB were found (Weninger et al., 1999), which revealed only a small effect (*g* = 0.30 [95CI −0.32, 0. 92], P = 0.34; Figure 7A). This supports the idea that CRH has only a modest effect on the ARDEB. The meta-analytic result may suffer from insufficient precision as the cumulative sample size was small (N = 20, 21). As publication bias appears to affect the literature, this small effect could be an overestimate.

### 3.12 Crhr1 gene knockout has a large effect on rodent anxiety

CRH exerts its biological action via two receptors known as CRHR1 and CRHR2. The two receptors are pharmacologically distinct and only the former has been widely studied in the context of anxiety (Owens and Nemeroff, 1991; Paez-Pereda et al., 2011). Meta-analysis (Gammie and Stevenson, 2006; Liebsch et al., 1999; Liebsch et al., 1995; Müller et al., 2003; Smith et al., 1998; Trimble et al., 2007) found that, in contrast to the *Crh* knockout, deletion of *Crhr1* had a large anxiolytic effect on ARDEB (*g* = −1.0 [95CI −1.30, −0.70], P = 6.64 × 10^−11^; Figure 7B). The discordance between *Crh* and *Crhr1* knockout effects has previously been attributed to the action of other peptide ligand(s) of *Crhr1*, either urocortin or another, unidentified ligand (A. J. Dunn and Swiergiel, 1999).

### 3.13 Species and sex differences in ARDEB

Rats and mice have differences in their defensive behavior when exposed to predators or predator cues (Blanchard, 2001). We used the synthetic data to examine species differences in baseline ARDEB prior to anxiotropic manipulation. The LD box showed the most substantial difference between species −6.88 % [−13.3; −0.46], P = 0.04, but in general, inter-species differences in naive ARDEB were minor when compared to the overall variance (Figure 8A).

Where rat data were available, inter-species differences between the effects of anxiety-related interventions were investigated. There were only minor inter-species differences in the meta-analytic effect sizes of the interventions (Figure 8B). A striking exception to this trend was a large rat-mouse difference in the response to restraint: this treatment appeared strongly anxiogenic for rats, but modestly anxiolytic for mice.

We investigated sex differences in ARDEB, but found that the anxiety studies contained ~18× fewer experiments on female rodents than males, rendering any meta-analytic estimates of female ARDEB imprecise.

### 3.14 Validity of the ARDEB assays

Of the interventions analyzed above, only diazepam has extensive clinical evidence supporting its ability to alter human anxiety. The large anxiolytic diazepam effect size observed with the ARDEB assays verifies their validity (Figure 9A). The stressors—isolation, acute pain and restraint—would all be expected to produce increases in human anxiety, and all show anxiogenic effects in the ARDEB, thus also verifying the validity of these assays (Figure 9A), with the exception of the surprisingly small social isolation effect (0.21 *g*). Establishing the validity of animal models relies partly on showing concordance between models (Campbell and Fiske, 1959; van der Staay, 2006). To explore the concordance between the three assays, we conducted regression analyses on all possible two-way comparisons of ΔARDEB (Figure 9B- Figure 9D). The LD-EPM and OF-LD comparisons of ARDEB changes both showed 60% concordance, supporting the idea that the three assays were measuring similar aspects. However, surprisingly, the OF-EPM comparison revealed that the two methods were discordant (R^2^_adj_ = −0.01, Figure 9D).

## 4 Discussion

### 4.1 Summary of evidence

Inspection of the forest plots reveals that all of the primary publication sets include experimental effect sizes that are discordant, either in direction (anxiolytic versus anxiogenic) and/or magnitude. The generality of discordance in the literature emphasizes the utility of metaanalysis to behavioral neuroscience to give a quantitative overview and to synthesize the best evidence available. Of ten analyses of putative anxiotropic interventions, eight yielded at least moderate meta-analytic effect sizes and two produced small effect sizes (Figure 9). The synthetic data strongly confirm that diazepam, the serotonergic system, environmental stressors, and *Crhrl* influence an anxiety-like process in the mouse brain.

### 4.2 Limitations

This study is limited by its exclusive use of English-language published data. Some studies had to be excluded from the meta-analysis during the full text scan because they did not report measures of variance. Only studies that reported time or percent time spent in exposed arena could be selected for meta-analysis. We found no knowledge gaps *per se*, as all ten proposed anxiety-related factors had at least two studies.

Nevertheless, *Htt* overexpression, *Crh* knockouts and the non-anxiety genes had limited cumulative sample sizes (N_cumulative_ < 64, 64). Of the six factors for which publication bias was examined, three were affected. The presence of publication bias in the larger data sets suggests that inclusion of further data to the smaller meta-analyses would be expected, on average, to lower these effect sizes as well. Heterogeneity was at least moderate (I^2^ > 50%) in five of the meta-analyses, indicating that the random effects model is insufficient to explain the variance in these data sets. Thus, laboratory, strain, assay type and other protocol variations played variable roles across factors. Heterogeneity could in theory be reduced by increased standardization (Crabbe et al., 1999). Multilevel regression models of these data may be able to account for the unexplained variance (Yildizoglu et al.,2015).

### 4.3 Assay validity

The validity of each ARDEB assay was originally tested with a panel of anxiotropic agents (Crawley and Goodwin, 1980; Pellow et al., 1985; Simon et al., 1994). In the decades since the variability of assay results and the disappointing clinical outcome of compounds identified with these preclinical assays raise new questions about their validity (Griebel and Holmes, 2013). The diazepam, restraint and acute pain synthetic data shown here support the ARDEB assays’ validity, though two other results raise doubts: (1) the social isolation effect on ARDEB is weaker than expected (Figure 9A); (2) the failure of EPM and OF to reproduce each other’s outcomes (Figure 9D). The EPM-OF metaregression discordance is an exploratory observation that could be verified with a formal method comparison with animals run through all three assays (Bland and Altman, 1999).

How might the validity of the ARDEB assays be tested further? First, meta-analyses of additional known anxiotropic agents will help assess the assays’ strengths and weaknesses. Second, assay validity assessment would be helped by researchers making their video or tracking data available (ideally with experimental metadata in a standard file format), similar to data sharing efforts currently underway in neurophysiology. Anxiety assay validity may also be tested with new instrumentation that allows the estimation of animal pose (Nanjappa et al., 2015; Wiltschko et al., 2015) and that will make complex, ethological relevant anxiety assays (Blanchard et al., 2001) increasingly accessible for routine analysis (Schaefer and Claridge-Chang, 2012). Looking backward (meta-analysis and data sharing) and forward (more refined anxiety assays) are both valuable to rodent anxiety research. It has been suggested that research consortia should form to overcome the cost restrictions of large rodent sample sizes (Button et al., 2013); perhaps a consortium could form around the problem of anxiety assay validation.

### 4.4 Disconnect between htr1a & crhr1 preclinical results and clinical efforts

Meta-analysis of *Htr1a* overexpression revealed it has a moderate anxiotropic effect (−0.6 *g*), smaller than the bias-corrected diazepam effect (−0.85 *g*), suggesting that compounds aiming to increase 5-HT1A function may be a poor strategy to reduce anxiety. This view is supported by clinical meta-analyses that have concluded that drugs targeting 5- HT1A—the azapirones—appear inferior to benzodiazepines for generalized anxiety disorder (Chessick et al., 2006) and that there is insufficient evidence to support azapirone use in panic disorder (Imai et al.,2014). It appears that clinical adoption of the azapirones was/is not informed by the preclinical genetic evidence base. A second type of preclinical-clinical disconnect is observed with the *Crhr1* knockouts. The synthetic preclinical data indicate that *Crhr1* knockout produces a very large reduction of rodent anxiety (*g* = −1.0 [95Ci −0.7, −1.3], I^2^ = 13%, N_cumulative_ = 105, 99). However, at least one clinical trial of a CRHR1 antagonist for generalized anxiety disorder showed no benefit over placebo (Coric et al., 2010). The discrepancy between the efficacy of *Crhrl* knockouts and inefficacy of CRHR1 antagonists in patients remains unexplained.

### 4.5 A paradox in HTT-SSRI anxiety effects

Drugs that inhibit SERT, the SSRIs, are recommended as the first line of pharmacological treatment for anxiety (Baldwin et al., 2014). Blocking SERT-mediated reuptake of serotonin from the synaptic cleft is the proposed mechanism of SSRI anxiety reduction, although rodent studies of chronic SSRI effects on ARDEB have been inconclusive (Griebel and Holmes, 2013; Perez-Caballero et al., 2014). Given the inhibitors’ clinical use, it is paradoxical that *Htt* knockouts have elevated anxiety relative to controls (0.57 *g*) and that *Htt* overexpression dramatically reduces rodent anxiety (−0.94 *g*). The reason for this drug/gene incongruence is not clear. In some cases, the authors of the primary *Htt* knockout studies have not discussed it (Carroll et al., 2007; Kalueff et al., 2007a; Moya et al., 2011; Schipper et al., 2011). Other authors have remarked that the underlying reason remains unclear (Holmes et al., 2003b; Lira et al., 2003) or have called the validity of ARDEB assays into doubt (Pang et al., 2011). As both genetic knockouts and SSRIs are expected to produce monotonic, systemic reductions of SERT function, this incongruence is not easily explained by models of serotonin conflicting action that invoke distinct 5-HT circuits in the brain with opposing effects on defense (Deakin and Graeff, 1991). Others have proposed two explanatory hypotheses. The first is that increased anxiety arises from developmental alterations present in *Htt* knockouts not present in chronically drug-treated animals (Holmes et al., 2003b; Olivier et al., 2008; Zhao et al., 2006). This hypothesis could be tested with conditional knockdown models, i.e. in animals with *Htt* only deleted at the adult stage. While systematic review of PubMed and EMBASE did not identify any published reports of post-developmental *Htt* knockout experiments (e.g., using floxed *Htt*), researchers have analyzed the anxiety-related effects of conditionally ablating the *Pet-1* gene. *Pet-1* is a transcription factor with an expression range that overlaps closely with the expression of *Htt*. In mice with *Pet-1* removed in adulthood, mRNA levels of *Htt* are substantially reduced (Chen Liu et al., 2010). Like *Htt* knockouts, these mice show increased anxiety-like behaviors in multiple ARDEB assays (Chen Liu et al., 2010), eroding confidence in the developmental alteration hypothesis. A second hypothesis to explain the *Htt*/SSRI paradox is that there is a J-shaped relationship between *Htt* function and anxiety, i.e., both wild-type and knockout animals would have higher anxiety relative to animals with intermediate function (Olivier et al., 2008). The SSRI/knockout paradox is also observed in depression assays, though interfering RNA knockdown of *Htt* in adult mice reduced the forced swim test measure of depression (N = 10, 10) (Thakker et al., 2005).

## 5 Conclusions

This study confirms that diazepam, two environmental stressors and three genes influence rodent anxiety as measured by defense behavior assays. These anxiety-related interventions (diazepam, *Htr1a* gene knockout, *Htt* gene knockout, *Htt* gene overexpression, acute pain, restraint and *Crhr1* gene knockout) can be used as reference manipulations when establishing other anxiety models. The rodent anxiety literature is affected by publication bias that amplifies effect sizes. For the panel of ten interventions, there is strong EPM-LD and OF-LD ARDEB assay concordance, but EPM and OF did not reproduce each other. The meta-analytic results bring several preclinical-clinical incongruencies into sharp relief: the weakness of *Htr1a* overexpression contrasting with the clinical use of azapirones, the potently anxiogenic *Crhr1* knockout contrasting with the clinical failure of CRHR1 antagonists, and the anxiogenic SERT knockout contrasting with the clinical use of SSRIs as anxiolytic drugs. Meta-analysis has the ability to aggregate information and resolve discordance in the primary literature, something of particularly use to behavioral neuroscience where most primary articles describe experiments with poor precision (Button et al., 2013). Precise estimation of effect magnitudes (Claridge-Chang and Assam, 2016) is important both to better understand animal model strengths/weaknesses and to improve the ability of preclinical studies to guide clinical investigation. The formation of multi-lab consortia to coordinate the examination of important hypothesized anxiety factors would be one promising way to increase the reliability of rodent anxiety data (Button et al., 2013). New, automated methods of behavioral imaging will also play a role in better preclinical models (Schaefer and Claridge-Chang, 2012). Another possibility would be to use small animal models (worms, flies, and zebrafish) that allow large sample sizes and powerful genetic tools to complement rodent experiments (Mohammad et al., 2016).

## Supplementary legends

**Supplementary Table 1. Characteristics of included experiments.** Characteristics of experiments from included studies are listed with Pubmed ID, year of study, and figure panel. The assay type, assay duration, variable used in experiment, route of injection, drug dosage, treatment duration, species, strain and gender are also detailed. Assay duration and treatment duration are listed in minutes. Dosage is listed in mg per kg body weight of animal. Cells containing NS (Not Specified) indicate that the information was not available in the study.

**Supplementary Information 1.** Spreadsheet containing extracted data (.xlsx file). Each dataset is in a separate sheet in the Excel file.

**Supplementary Information 2.** Extracted meta-analytic data in R-compatible format (.RData file)

**Supplementary Information 3.** R markdown code (.Rmd) used for meta-analysis and plotting

## 7 Author contributions

Conceptualization, FM and ACC; Software Programming, JH and FM; Investigation, FM, JH, CLL, JHW, DJJP and BL; Validation, FM, JH, CLL, JHW, DJJP and BL; Data Curation, FM; Writing – Original Draft, FM, JH and ACC; Writing – Review & Editing, ACC; Visualization, FM, JH and ACC; Supervision, ACC; Project Administration, ACC; Funding Acquisition, ACC. All authors have agreed to the final content.

## Funding information

The authors were supported by Biomedical Research Council block grants to the Neuroscience Research Partnership and the Institute of Molecular and Cell Biology. FM and ACC also received support from Duke-NUS Graduate Medical School. JH received support from the A^*^STAR Graduate Academy. ACC received additional support from a Nuffield Department of Medicine Fellowship, a Wellcome Trust block grant to the University of Oxford, A^*^STAR Joint Council Office grant 1431AFG120 and NARSAD Young Investigator Award 17741. The funders had no role in study design, data collection and analysis, decision to publish, or preparation of the manuscript.

